# The impact of struvite and ash recycling-derived fertilizers on microbial phosphorus mobilization capabilities and community structure in a *Lolium perenne* field trial

**DOI:** 10.1101/2024.01.16.575387

**Authors:** Lea Deinert, SM Ashekuzzaman, Patrick Forrestal, Achim Schmalenberger

## Abstract

Rock phosphate is a non-renewable primary source for mineral phosphorus (P) fertilizers that intensive agriculture is highly dependent on. To avoid P fertilizer shortages and limit negative environmental impacts, circular economy approaches are needed with recycling-derived fertilizer (RDF) applications such as struvites and ashese. Hence, a grassland field trial was conducted with four RDFs, two struvites (potato wastewater, municipal wastewater) and two ashes (poultry-litter ash, sewage-sludge ash) at a P application rate of 40 kg P ha^-1^ (n=5). The impact of the RDFs on the soil microbial P cycling community was compared to conventional mineral P-fertiliser and a P-free control. Topsoil samples were taken directly after *Lolium perenne* grass cuts at months three, fife and 15. Cultivable phosphonate and phytate utilizing bacteria, potential acid and alkaline phosphomonoesterase activity, and *phoC* and *phoD* copy numbers responded stronger to seasonal effects than treatment effects. No significant overall effect of the fertilizer application was detected in the beta diversity of the bacterial and fungal communities after 15 months, but individual phylogenetic groups were affected by the treatments. The ash treatments demonstrated some distinguishing results across the bacterial and fungal community, with significantly higher relative abundance of Firmicutes and Rokubacteria and lower relative abundance of Actinobacteriota. Sewage-sludge ash had significantly lowest abundances of genera *Bacillus* and *Bradyrhizobium* (month 15s) that are well known for their P cycling abilities. The struvite RDFs either positively influenced the P cycling microbial community or did not affect it at all, while demonstrating better tri-calcium phosphate solubilizing capabilities after the 3 months harvest. These findings indicate that struvites could be a suitable replacement for conventional P fertilizers in the future.

## 1. Introduction

Phosphorus (P) is a scarce and non-renewable resource. Still, over 80% of the P demand for agricultural purposes is covered by fossil P resources to produce linear economy mineral P fertilizers, less than 20% of P is sourced through the circular economy (Chowdhury *et al*., 2014). Efforts to increase the P recycling rate are essential to secure feeding the growing world population in the future (Nations, 2019). In natural soils, plant available P replenishment from the soil P pool in the form of orthophosphate occurs as weathering from apatite, a primary P mineral. This ultimately leads to the slow release of orthophosphate into the soil solution for plant uptake (Plante, 2007). Plant available orthophosphate is immobilized quickly in soil via adsorption to particles and clay, precipitation into inorganic calcium, aluminium or iron salts or integration into organic P compounds. Therefore, reliance on this weathering process is not feasible for crop production, because the P removal via crop harvests far outstrips plant available P replenishment (Barea & Richardson, 2015). Soil microbes play an important role in the mediation of soil P. While water-soluble, mineral P fertilizer quickly immobilizes after soil application, soil bacteria such as actinomycetes and saprophytic fungi are constantly involved in P mobilization and immobilization reactions (Richardson & Simpson, 2011).

Application of readily available mineral P in conventional fertilizers can disturb plant-microbe networks and the microbial P cycling activities as well as impair natural P regulation ability of the soil in the long term (Huang *et al*., 2019). Then, the dependency on fertilizer input for efficient plant growth increases even further (Bünemann *et al*., 2011). Over 70% of agricultural soils are not P deficient in general, they rather only contain P predominantly in plant inaccessible forms (Oburger *et al*., 2011). This is due to the phosphorus sorption capacity (PSC) of the soil. The PSC varies with soil type due to different characteristics (pH, clay and organic matter content, aluminium, calcium and iron concentrations). Continuation of mineral P fertilizer application to improve crop cultivation despite high PSC causes P accumulation in the soil, until a certain degree of phosphorus saturation (DPS) is reached, making the soil susceptible to leaching and P run-off (Werner & Wodsak, 1995, Casson *et al*., 2006). The uncontrolled loss of nutrients causes eutrophication of water bodies with detrimental effects for aquatic life (Correll, 1998, Torrent *et al*., 2007, Ulén *et al*., 2007). Already a DPS value of 25% or above has been determined as critical, where the amount of P leached from agricultural soil becomes inacceptable (Breeuwsma *et al*., 1995). Therefore, it is important to maintain microbial P cycling capacities in agricultural soil for the mobilization of P from plant inaccessible P pools (Oehl *et al*., 2001).

Phosphorus demands in grasslands to 2050 are projected to be in the region of 4 to 12 million tons of P per year (Mogollón *et al*., 2018). This demand is not sustainable if met with the declining fossil P reserves alone. Therefore, it is crucial to develop and evaluate recycling-derived fertilizers produced via various nutrient recovery technologies from waste streams to substitute fossil P use as much as possible (MacDonald *et al*., 2011). There are other approaches already in place for the improvement of nutrient re-use, such as direct application of manure and sewage sludge to fields (Huygens & Saveyn, 2018). These practices however are associated with issues such as high concentration of pollutants, like pathogens, endocrine disruptors and toxic heavy metals, which pose threats to food security and also impact the soil microbial biomass (Alvarenga *et al*., 2016, Charlton *et al*., 2016). Recycling-derived fertilizers that stem from processed waste products may be a promising alternative to substitute agricultural P demand instead of the use of raw waste products.

Currently, there is a lack of understanding how recycling-derived fertilizers such as ashes and struvites derived from various feedstocks affect plant P availability and, in particular, how they affect the soil P mobilizing microbiota, which is responsible for the mobilization of plant unavailable P in natural soil environments. Soil microorganisms are an essential actor in the P turnover and are involved in all key transformations of the soil P cycle, mineralization and immobilization, dissolution and precipitation and sorption and desorption (Bünemann *et al*., 2011). Their involvement in the uptake, release and redistribution of P significantly influences plant growth performance (Gyaneshwar *et al*., 2002).

The objective of this study was to evaluate the performance of four RDFs for their impact on the P-mobilizing soil microbiota in terms of phosphonate and phytate utilization, tri-calcium phosphate solubilization, potential phosphatase activity, P cycling gene copy numbers and abundance, and shifts in the bacterial and fungal community structure. Potato processing wastewater struvite (PWS), municipal wastewater struvite (MWS), poultry litter ash (PLA) and municipal sewage sludge ash (SSA) were applied in a single application of 40 kg P ha^-1^ which is in accordance with agronomic advise for a P-deficient soil. It was hypothesized that the RDFs promote plant growth in a similar manner compared to superphosphate (SP) fertilizer but are potentially less impactful on the soil microbial P cycling processes.

## 2. Materials and Methods

### 2.1 Description of Field Trial Layout and Sampling Procedure

The field trial was conducted at Teagasc Johnstown Castle in Wexford, Ireland (N52°17’47”, W6°30’29”). The sandy loamy soil had an initial Morgan’s P concentration of 1.3 mg P L^-1^, index 1 P-deficient by the Irish system, with a pH of 5.6 (Karpinska *et al*., 2021). The amount of P fertilizer used for the treatments was 40 kg P ha^-1^. The RDFs PLA, SSA, MWS and PWS were applied to plots (6 m x 2 m) in five replicates in a randomised complete block design to balance gradients and natural variability in the field. Furthermore, a control without P(0 kg P ha^-1^) was included. All plots received an equivalent of 125 kg ha^-1^ of Nitrogen in the form of Calcium Ammonium Nitrate, 155 kg ha^-1^ of Potassium and 20 kg ha^-1^ of Sulphur during the first application. Then, these were applied at N (100 kg ha^−1^), K (75 kg ha^−1^) and S (20 kg ha^−1^) for second and third applications, respectively, for the remaining grass growing season over a year. All RDF fertilizers were applied by hand in April 2019, to ensure equal distribution within the plots and avoiding contamination between different plots. The area surrounding the plots was treated with fluroxypyr, an auxin-type herbicide used to control annual and perennial broadleaf weeds (Washington State Department of Transportation, 2017), to suppress grass growth around the plots.

Directly after the grass cuts performed in May (harvest, after one month), July (harvest, after 3 months), September (harvest, after 5 months) and July 2020 (harvest, after 15 months) soil samples were taken with a soil corer from the topsoil (10 cm) in a W shaped pattern.

The soil samples of the treatments SP0, SP40, PLA40, SSA40, MWS40 and PWS40 were sieved on-site (mesh size 3.35 mm). The samples were taken aseptically by disinfecting the sampler and sieves with isopropyl alcohol (60 % v/v) between taking each sample. The samples were stored on ice for transport and subsequently stored at −20°C for DNA extraction and for the remaining experiments at 4°C. Within 24 hours of the harvest, MPN and CFU analysis was performed.

### 2.2 Sample Processing and Analysis

The soil pH, Most Probable Number (MPN) and Colony Forming Unit (CFU) analysis, alkaline phosphatase ACP and acid phosphatase ALP enzyme activity assays, *phoC* and *phoD* qPCR and *phoD* functional amplicon sequencing were performed as described recently in detail (Deinert *et al*., 2023). In brief, pH was measured with 5 g of air-dried soil in 20 mL of 0.01 M CaCl_2_ solution. MPN determination was carried out via serial dilution of samples (1 g soil in 10 mL sterile rotated at 75 rpm for 30 min at 4°C). Microtiter plates contained MM2PAA or MM2Phy medium with phosphonate or phytate as sole P source, or R2 medium (Reasoner & Geldreich, 1985, Fox *et al*., 2014) were inoculated with 20 µL of the serial dilution (technical replicates of 5) and incubated at 25°C for 14 days to obtain MPN values g^-1^ (Blodgett, 2023) Tri-calcium-phosphate (TCP) solubilizing CFUs were obtained via plating of 100 μL of serial dilutions (Fox *et al*., 2014) in three technical replicates (incubated at 25°C, 14 d). The cultivation of soil microbes on the solid medium supplemented with tri-calcium phosphate was analysed by counting the colonies which formed a clearing zone indicating the solubilization of the TCP in the agar. Measuring the potential acid and alkaline phosphomonoesterase activity in soil was conducted following standard protocols (Tabatabai & Bremner, 1969) as described recently (Deinert *et al*., 2023). The available P was measured using the Morgan’s soil P test, which is typically employed for examining Irish soil P status (Daly & Casey, 2005) as previously detailed (Deinert *et al*., 2023).

DNA extraction was performed using the DNeasy PowerSoil Pro kit (QIAGEN GmbH, Hilden, Germany) according to the manufacturer’s instructions and quantified using the Qubit Fluorometer (dsDNA HS assay kit, Life Technologies, Carlsbad, CA, USA).

ITS PCR-DGGE was performed in a nested PCR approach as described elsewhere (Schmalenberger & Noll, 2014). For the first PCR the primers ITS1 F (5‘-CTT GGT CAT TTA GAG GAA GTA A-3‘) and ITS 4 (5’-TCC TCC GCT TAT TGA TAT GC-3’) (White *et al*., 1990) were utilized to primarily amplify *Ascomycota*, *Basidiomycota* and *Zygomycota*. Cycling conditions were as follows: denaturation at 95 °C (4 min), followed by 30 cycles of denaturation (94 °C, 45 s), annealing (50 °C, 45 s), extension (72 °C, 60 s) and a final elongation (72 °C for 10 min). PCR products was diluted 1:10 and used as a template for the second PCR with primers ITS1 F-GC (5’-CGC CCG CCG CGC GCG GCG GGC GGG GCG GGG GCA CGG GGG GCT TGG TCA TTT AGA GGA AGT AA-3’) and ITS 2 (5‘-GCT GCG TTC TTC ATC GAT GC-3‘) (White *et al*., 1990). Cycling conditions were as above but broken up into two rounds of 15 cycles with 60 °C annealing and followed with 20 cycles with annealing at 50 °C; a final extension at 72 °C for 5 min completed the cycling. The DGGE conditions were the same as described before (Schmalenberger & Noll, 2014) albeit the urea/formamide denaturing gradient was adjusted to 30-50 %.

16S and ITS Illumina next-generation sequencing of soil DNA extracts originating from the 15 months soil sampling was performed by Novogene (UK) Company Limited using genomic DNA from the same extractions as described above. To analyse the 16S rRNA gene, the primers for the V4-V5 region (515F: 5’-GTG CCA GCM GCC GCG GTA A-3’, 907R: 5’-CCG TCA ATT CCT TTG AGT TT-3’), and for the ITS analysis the primers for ITS1 (ITS5-1737F: 5’-GGA AGT AAA AGT CGT AAC AAG G-3’, ITS2-2043R: 5’-GCT GCG TTC TTC ATC GAT GC-3’) were chosen with barcodes for PCR amplification using Phusion^®^ High-Fidelity PCR Master Mix (New England Biolabs, Ipswich, MA, USA). The PCR products were then mixed at equal density ratios and purified with a Gel extraction kit (QIAGEN, Düsseldorf, Germany). Sequencing libraries were generated with the NEBnext^®^ Ultra^TM^ DNA Library Prep kit for Illumina^®^ (New England Biolabs, Inc., Ipswich, MA, USA). After quality assessment via Qubit (Life Technologies, Carlsbad, CA, USA) and qPCR, the libraries were sequenced using a NovaSeq PE250 platform to generate 250 bp paired end reads.

Sequencing of *phoD* was carried out at the University of Minnesota Genomics Centre (UMGC, MN, USA) using PE300 MiSeq amplicon sequencing as described previously (Deinert *et al*., 2023). Briefly, *phoD* gene fragments were amplified in triplicates with primers *phoD*-F733 (5’-TGG GAY GAT CAY GAR GT-3’) and *phoD*-R1083 (5’-CTG SGC SAK SAC RTT CCA-3’) (Ragot *et al*., 2015) in 25 µL reactions with KAPA2G Robust HotStart PCR kit (KAPA Biosystems, Roche; 0.5 U KAPA2G Robust HotStart DNA Polymerase, 1x KAPA2G Buffer A, 0.5 mM MgCl_2_, 0.2 mM dNTPs) and 1 M betaine, 0.8 mM of each primer and 0.5 µL DNA template. Cycling conditions were 3 min of initial denaturation (95°C), 35 cycles of denaturation (95°C, 30 s), annealing (58°C, 15 s), elongation (72°C, 15 s), and a final extension (72°C, 3 min). PCR products were purified with the GenElute PCR Clean-up kit (Sigma Aldrich, St. Louis, MO, USA) as per instruction manual and triplicates were pooled and diluted 1:5 for a second PCR with Illumina adapter sequences. Here, the KAPA HiFi HotStart Readymix (KAPA Biosystems, Roche, 0.5 mM MgCl_2_) was used in 25 µL reactions with 0.4 mM of each primer and 1 µL template. Cycling conditions for initial denaturation were the same as above, followed by 15 cycles of denaturation (95°C, 20 s), annealing (65°C, 15 s), elongation (72°C for 15 s), and a final extension (72°C, 1 min). Technical replicates were pooled, purified (GenElute PCR Clean-up kit, Sigma Aldrich) and quantified using a Qubit Fluorometer (Life Technologies, dsDNA HS assay kit). Samples were shipped to UMGC on dry ice for library preparation, sequencing and demultiplexing.

*phoC* qPCR was conducted with the primers *phoC*-A-F1 (5‘-CGG CTC CTA TCC GTC CGG-3‘) and *phoC*-A-R1 (5’-CAA CAT CGC TTT GCC AGT G-3’) (Fraser *et al*., 2017) with a Roche LightCycler^®^ 96 (Roche Diagnostics, Mannheim, Germany) using 5 µL KAPA SYBR FAST 2x qPCR Master Mix (KAPA Biosystems, Wilmington, MA), 0.3 mM of each primer and 1 µL of DNA template (20 ng µL^-1^). The cycling conditions were initial denaturation (95 °C, 5 min), followed by 45 cycles of denaturation (95 °C for 3 s), annealing (60 °C for 20 s), elongation and detection (72 °C, 10 s) followed by a melting curve. *phoD* qPCR was carried out as described previously (Graça *et al*., 2021) with primers ALPS-F730 (5’-CAG TGG GAC GAC CAC GAG GT-3’) and ALPS-R1101 (5’-GAG GCC GAT CGG CAT GTC G-3’) (Sakurai *et al*., 2008) and the same cycling conditions as for *phoC*. Absolute quantifications were reported in copies per g^-1^ soil.

### 2.3 Statistical Analyses

The soil potential phosphomonoesterase activity assay, MPN and CFU analysis, *phoD* and *phoC* gene quantification via qPCR, were analysed for statistical significance in SPSS (IBM, Version 26). Normality was assessed via the Shapiro-Wilk test. The homogeneity of variance was evaluated using Levene’s test and if both assumptions were fulfilled, a one-way analysis of variance (ANOVA) was performed, applying the Tukey HSD post-hoc test for pairwise comparison of the treatments (P<0.05) to assess significant differences between treatment means. Data violating one or both assumptions of normality and homogeneity of variance were transformed either using logarithm or square root transformation, the ANOVA was repeated, and the untransformed results were reported. When homogeneity of variance was not achieved, a Games-Howell post-hoc analysis was chosen. For data violating the normality assumption even after transformation, the non-parametric Kruskal-Wallis test was performed. Analysis of the band pattern of the DGGE analyses was conducted with Phoretix 1D software (Totallab, Newcastle, UK). The resulting binary matrix was used in a canonical correlation analysis (CCA), testing effects of measured environmental factors and visualizing the CCA biplot using the R packages vegan, mabund, permute and lattice, applying a permutational multivariate analysis of variance (PERMANOVA) test using 999 permutations.

Demultiplexed, paired-end 16S rRNA (bacterial) and ITS (fungal) sequence reads obtained from Novogene UK Co. Ltd (Cambridge, UK) were imported into QIIME2 2020.8 (Bolyen *et al*., 2019). The paired-end reads were joined, quality filtered and denoised via the q2-dada2 plugin (Callahan *et al*., 2016). The resulting amplicon sequence variants (ASV) were aligned using the q2-alignment with mafft (Katoh & Standley, 2013) and a phylogenetic tree was built via q2-phylogeny with fasttree2 (Price *et al*., 2010). Samples were rarefied to 29431 and 61307 sequences per sample for the 16S and ITS analysis respectively, and the alpha diversity metrics (observed species and Faith’s Phylogenetic diversity (Faith, 1992)) and beta diversity metrics (Bray-Curtis dissimilarity, Jaccard distance, unweighted UniFrac (Lozupone & Knight, 2005) and weighted UniFrac (Lozupone *et al*., 2007) were assessed using the q2-diversity plugin. Taxonomy assignment was accomplished using the q2-feature-classifier (Bokulich *et al*., 2018), training with a SILVA 13.8 99% reference data set for 16S rRNA (Quast *et al*., 2013, Yilmaz *et al*., 2014). The UNITE database was used for the fungal NGS analysis (Abarenkov *et al*., 2021). First, the reads in the reference data set were trimmed to the region of interest using the primers that had been used in PCR amplification prior to sequencing. Then, a Naïve Bayes classifier was trained with the trimmed reference sequences and their reference taxonomies. The mvabund and vegan packages in the R software (Version 4.0.3) were used to perform a CCA analysis based on the ASV table obtained during the QIIME2 analysis. Additionally, taxa bar plots were created to disclose the percentage relative abundance distribution of prevalent taxa among the treatments.

Demultiplexed *phoD* sequences (for details, see (Deinert *et al*., 2023)) were cleared of primers and poor-quality sequences (cutadapt software (Martin, 2011)). Paired end reads were merged (Usearch (Edgar, 2010)) and filtered (fastq_filter). Duplicate sequences were removed (fastx_uniques) and the UPARSE pipeline was used to cluster sequences into centroid operational taxonomic units (OTUs) (Edgar, 2013) at 75% sequence similarity (Fraser *et al*., 2015). Fungene was used to select a *phoD* reference database and to assign taxonomies (Fish *et al*., 2013). The mvabund and vegan packages (R software, Version 4.0.3) were used to create a CCA biplot and permutation analysis as well as taxa bar plots.

## 3. Results

### 3.1 Cultivation-Dependent Assessment of the P-Mobilizing and P-Solubilizing Bacteria

The results of the cultivation-dependent methods MPN and CFU for all three harvests are summarized in Table 1. For the harvest after 3 months, the phosphonate utilizing bacteria were significantly higher in the SP40 treatment when compared to SP0 and PWS40 but overlapped with the remaining treatments. That changed for harvests after 5 and 15 months, where no significant difference between treatments were observed. All phytate utilizing bacteria were statistically similar across all three harvests, though total numbers increased at month 15 for all treatments when compared to the two previous harvests. Cultivation in R2 medium also did not yield statistically different results for the treatments, albeit average numbers of heterotrophic bacteria were highest in SP40. Abundance of TCP-solubilizing bacteria in the SP40 treatment at month three were about ten-fold lower than for MWS40, SSA40 and PLA40, which was significant, while SP0 and PWS40 overlapped. This observation continued as a trend for the following two harvests, albeit no longer reached significance. Nevertheless, SP40 was had the second lowest (month 5) or lowest (month 15) abundance of TCP solubilizing bacteria.

**Table 1.** Overview of the bacterial P mobilization capabilities from phosphonoacetic acid (PAA) and phytate (Phy), and overall cultivation of heterotrophic bacteria in R2A medium analysed via the MPN approach, and tri-calcium phosphate (TCP) solubilization ability of the field trial harvests after 3, 5 and 15 months, different letters indicate significant difference (P < 0.05) within a column, ± represents standard error, n=5.

| Treatment | PAA (MPN g<sup>-1</sup> soil) |  | Phy (MPN g<sup>-1</sup> soil) |  | R2 (MPN g<sup>-1</sup> soil) |  | TCP (CFU g<sup>-1</sup> soil) |  |
| --- | --- | --- | --- | --- | --- | --- | --- | --- |
| Harvest after 3 Months |  |  |  |  |  |  |  |  |
| SP0 | 1.9E+06<sup>b</sup> | ±5.7E+05 | 3.7E+06<sup>a</sup> | ±7.1E+05 | 3.6E+07<sup>a</sup> | ±7.8E+06 | 4.7E+04<sup>ab</sup> | ±2.2E+04 |
| SP40 | 4.7E+06<sup>a</sup> | ±8.0E+05 | 3.8E+06<sup>a</sup> | ±1.2E+06 | 9.5E+07<sup>a</sup> | ±6.4E+07 | 6.7E+03<sup>b</sup> | ±6.7E+03 |
| PWS40 | 1.7E+06<sup>b</sup> | ±4.7E+05 | 3.5E+06<sup>a</sup> | ±7.4E+05 | 3.9E+07<sup>a</sup> | ±9.9E+06 | 6.0E+04<sup>ab</sup> | ±1.2E+04 |
| MWS40 | 6.3E+06<sup>ab</sup> | ±2.7E+06 | 5.7E+06<sup>a</sup> | ±1.4E+06 | 4.6E+07<sup>a</sup> | ±8.2E+06 | 7.7E+04<sup>a</sup> | ±1.9E+04 |
| PLA40 | 2.6E+06<sup>ab</sup> | ±7.4E+05 | 4.9E+06<sup>a</sup> | ±8.2E+05 | 5.6E+07<sup>a</sup> | ±1.7E+07 | 8.7E+04<sup>a</sup> | ±2.9E+04 |
| SSA40 | 2.6E+06<sup>ab</sup> | ±5.8E+05 | 4.9E+06<sup>a</sup> | ±1.8E+06 | 3.4E+07<sup>a</sup> | ±6.1E+06 | 1.2E+05<sup>a</sup> | ±4.4E+04 |
| Harvest after 5 Months |  |  |  |  |  |  |  |  |
| SP0 | 3.2E+06<sup>a</sup> | ±9.2E+05 | 7.0E+06<sup>a</sup> | ±2.4E+06 | n.a. |  | 6.7E+04<sup>ab</sup> | ±3.0E+04 |
| SP40 | 4.2E+06<sup>a</sup> | ±1.5E+06 | 3.4E+06<sup>a</sup> | ±6.6E+05 | 1.7E+09 | ±1.6E+09 | 2.0E+04<sup>ab</sup> | ±1.3E+04 |
| PWS40 | 3.8E+06<sup>a</sup> | ±1.1E+06 | 5.0E+06<sup>a</sup> | ±7.9E+05 | n.a. |  | 6.7E+03<sup>b</sup> | ±6.7E+03 |
| MWS40 | 2.9E+06<sup>a</sup> | ±1.0E+06 | 6.9E+06<sup>a</sup> | ±1.9E+06 | n.a. |  | 6.0E+04<sup>ab</sup> | ±6.0E+04 |
| PLA40 | 2.7E+06<sup>a</sup> | ±7.3E+05 | 7.0E+06<sup>a</sup> | ±2.5E+06 | n.a. |  | 8.7E+04<sup>ab</sup> | ±6.3E+04 |
| SSA40 | 1.6E+06<sup>a</sup> | ±3.7E+05 | 7.5E+06<sup>a</sup> | ±4.3E+06 | n.a. |  | 1.1E+05<sup>a</sup> | ±3.9E+04 |
| Harvest after 15 Months |  |  |  |  |  |  |  |  |
| SP0 | 2.0E+07<sup>a</sup> | ±4.5E+06 | 2.5E+07<sup>a</sup> | ±6.3E+06 | 1.0E+08<sup>a</sup> | ±3.0E+07 | 1.3E+04<sup>a</sup> | ±1.3E+04 |
| SP40 | 1.8E+07<sup>a</sup> | ±1.6E+06 | 1.6E+07<sup>a</sup> | ±2.3E+06 | 1.4E+08<sup>a</sup> | ±4.9E+07 | 6.7E+03<sup>a</sup> | ±6.7E+03 |
| PWS40 | 1.3E+07<sup>a</sup> | ±1.9E+06 | 2.6E+07<sup>a</sup> | ±5.9E+06 | 4.7E+07<sup>a</sup> | ±8.0E+06 | 2.0E+04<sup>a</sup> | ±1.3E+04 |
| MWS40 | 1.4E+07<sup>a</sup> | ±1.7E+06 | 2.0E+07<sup>a</sup> | ±2.8E+06 | 6.6E+07<sup>a</sup> | ±1.4E+07 | 4.7E+04<sup>a</sup> | ±1.7E+04 |
| PLA40 | 1.4E+07<sup>a</sup> | ±3.2E+06 | 2.8E+07<sup>a</sup> | ±3.0E+06 | 8.8E+07<sup>a</sup> | ±2.0E+07 | 2.7E+04<sup>a</sup> | ±1.9E+04 |
| SSA40 | 1.5E+07<sup>a</sup> | ±2.5E+06 | 2.3E+07<sup>a</sup> | ±4.5E+06 | 7.5E+07<sup>a</sup> | ±2.4E+07 | 3.3E+04<sup>a</sup> | ±2.1E+04 |

\*n.a. = not available

### 3.2 Phosphomonoesterase Activity and Gene Copy Number Assessment

At the time of harvest after 3 months, potential ACP activity was on average ten-fold higher than ALP activity. Treatments MWS40 and PLA40 showed a significantly higher potential ACP activity compared to mineral SP40, and the values for these were nearly twice as high as for the other treatments (Table 2). At the 5 months harvest, no statistically significant differences were found in the ACP activity for all treatments. However, the overall ACP activity was the highest at month 5, while ACP levels at months 3 and 15 were similar. Only the SSA40 treatment displayed significantly higher ACP activity compared to all other treatments at month 15, while all other treatments were similar (Table 2). ALP activity was not significantly affected by any of the treatments at any of the three harvests.

**Table 2.**
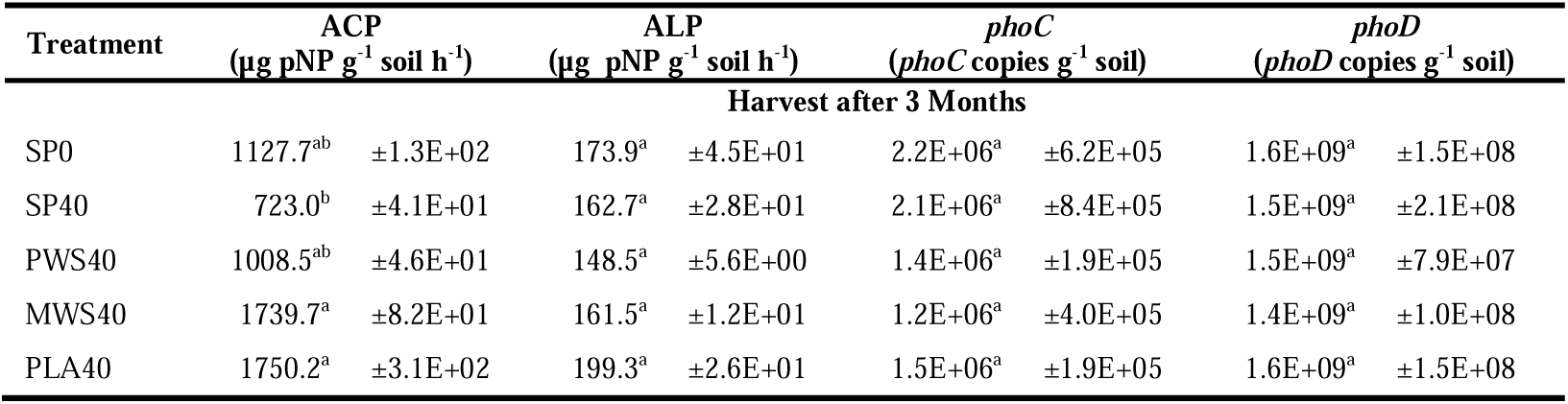
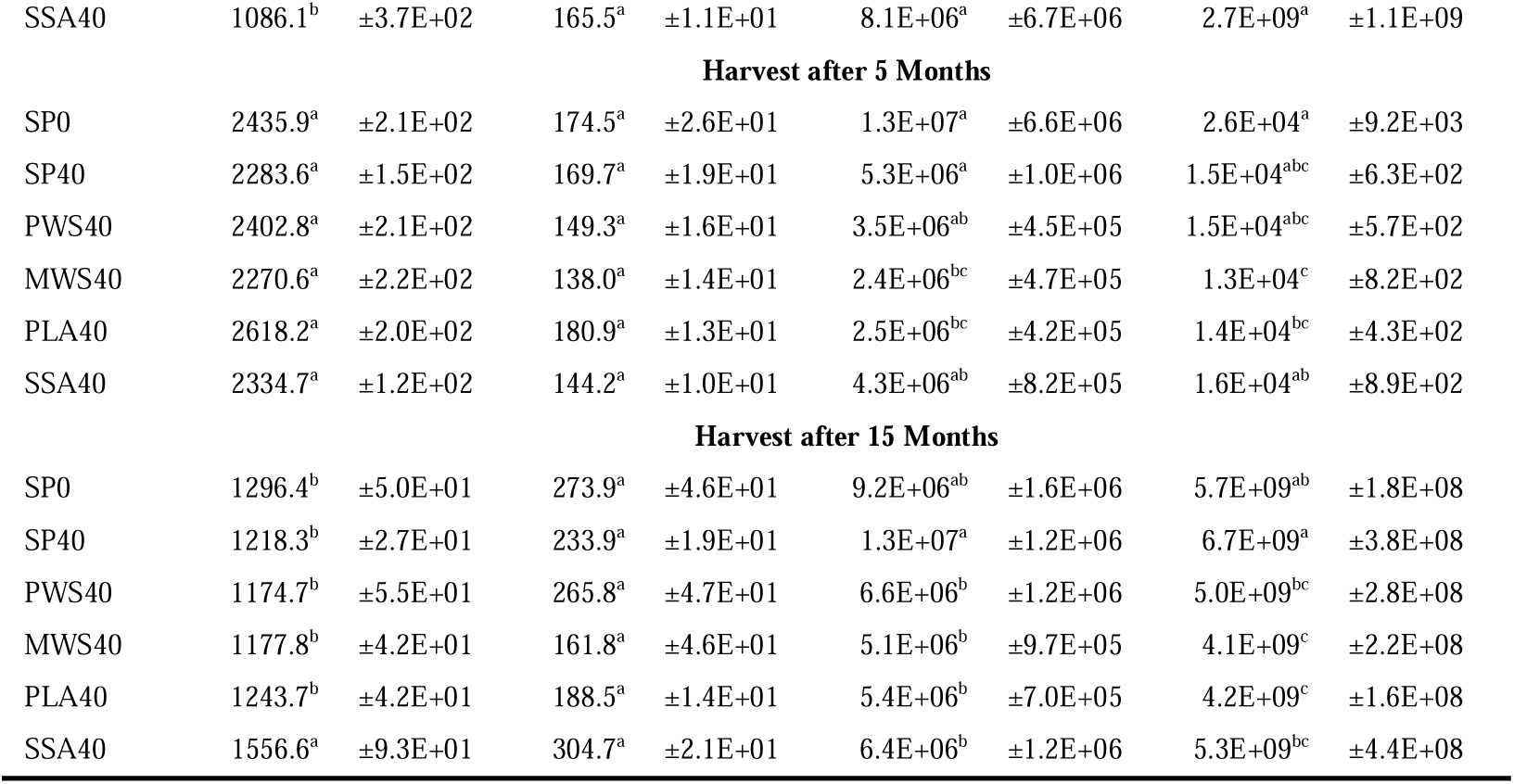
Collated data for the potential acid (ACP) and alkaline (ALP) phosphomonoesterase activity in field trial soil determined via spectrophotometry and phoC (acid) and phoD (alkaline phosphomonoesterase) copy numbers determined via qPCR for all three harvests (after 3, 5 and 15 months), different letters indicate significant difference (P < 0.05) within a column, ± represents standard error, n=5.

The *phoC* copy numbers were similar for all treatments after 3 months, but after 5 months the SP0 and SP40 treatments contained significantly higher *phoC* copy numbers compared to MWS40 and PLA40, while the other treatments overlapped. In the second year after 15 months, only the SP40 treatment still had significantly higher acid phosphomonoesterase copy numbers than the RDF treatments (Table 2). The *phoD* copy number values followed a similar trend as the *phoC* copy numbers. There was no statistically significant difference detected after 3 months between treatments but after 5 months the *phoD* copy numbers had dropped by several orders of magnitude. The copy numbers were observed to be the highest in the SP0 with significantly higher (P<0.05) values compared to MWS40 and PLA40, where MWS40 displayed the lowest *phoD* copy numbers with significantly lower values (P<0.05) than SP0 and SSA40, respectively. After 15 months harvest, the *phoD* copy numbers were at a comparable level to the 3-month harvest of the previous year, and a significantly higher (P<0.05) copy numbers were observed in the SP40 treatment compared to any of the RDF treatments. The lowest *phoD* copy number was found in the MWS40 treatment sample and this was significantly lower (P<0.05) compared to SP0 or SP40 treatments.

### 3.3 Bacterial and Fungal Community Analysis

2 461 486 single-end reads were obtained from the 16S rRNA bacterial community analysis of the DNA extracted from rhizosphere soil from the harvest after 15 months of the Teagasc field trial. After filtering, chimera removal and taxonomy assignment, Shannon alpha diversity ranged from 5.911 to 6.154, with the SP0 treatment having a significantly higher (P<0.05) alpha diversity than SSA40 (supplementary Table S1). Similarly, Fisher’s alpha was also significantly increased (P<0.05) in SP0 over SSA40. Other alpha diversity indices, such as Simpson, were not statistically different. The ten most relative abundant phyla out of 42, comprised over 95% of the total 16S rRNA ASVs, in descending order *Proteobacteria*, *Actinobacteria*, *Firmicutes*, *Acidobacteria*, *Planctomycetes*, *Verrucomicrobia*, *Chloroflexi*, *Gemmatimonadetes*, *Bacteroidetes* and *Nitrospirae* (Supplementary Figure S1).

Out of these phyla mentioned above, four were found to be differentially abundant, as shown in Figure 1. *Actinobacteriota* were significantly decreased (P<0.05) in PWS40 and PLA40 compared to the SP0 control treatment. Relative abundance of the phylum *Firmicutes* was significantly higher (P<0.05) in the PLA40 treatment compared to the SP40 and SSA40 treatment. Similarly, *Rokubacteria* displayed highest relative abundance in the PLA40 treatment as well, however it was only significantly different (P<0.05) compared to SP0. *Gemmatimonadetes* then were detected to significantly higher in relative abundance in SP40 than SP0 (P<0.05).

**Figure 1.**
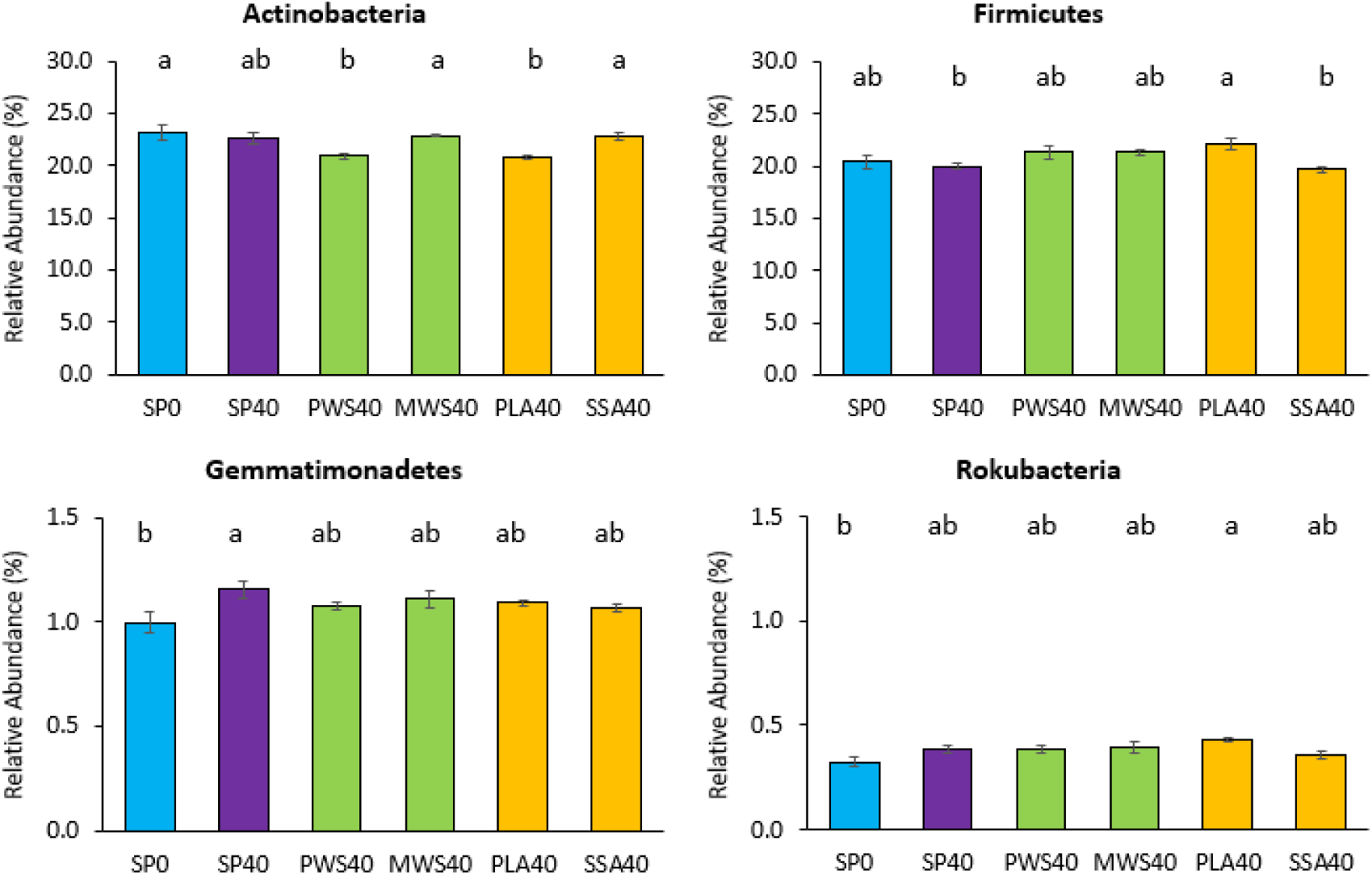
Mean relative abundance (%) of top 10 phyla of the harvest after 15 months of the 16S rRNA sequencing analysis that were differentially abundant, different letters indicate significant difference (P < 0.05), error bars represent standard error, n=5.

At genus level, only three out of the 29 most abundant genera (above 20% across all treatments) were significantly differentially (P<0.05) abundant (Figure 2). The genera *Bacillus* and *Bradyrhizobium* had both lowest abundance in the SSA40 treatment, the first was significantly lower (P<0.05) to the RDFs MWS40 and PLA40 and the latter to SP40. Similar to *Bradyrhizobium*, the genus *Streptomyces* was also highest abundant in SP40, however only significantly higher (P<0.05) compared to the PWS40 treatment.

**Figure 2.**
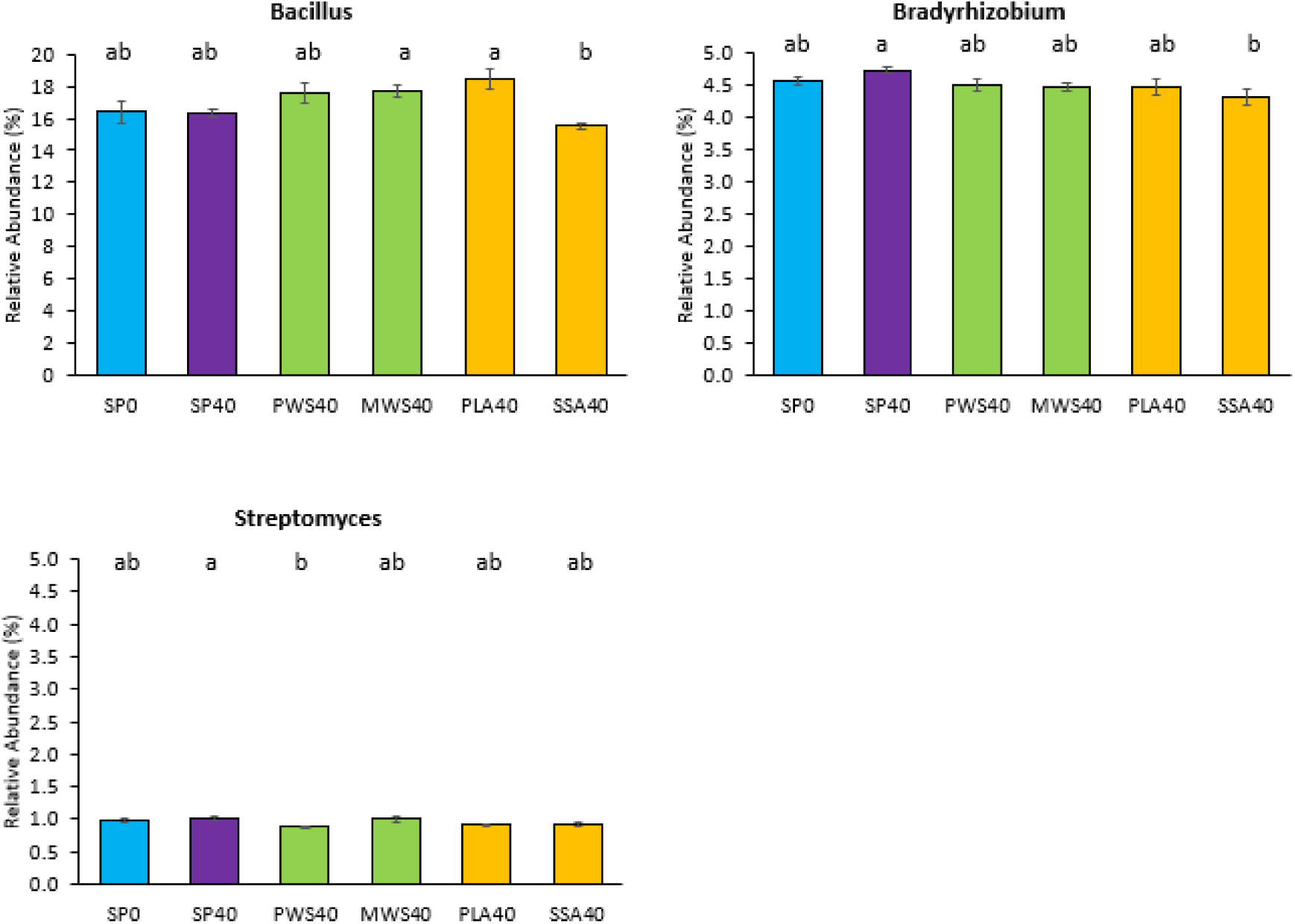
Mean relative abundance (%) of the genera after 15 months, that were statistically differential abundant, different letters indicate significant difference (P < 0.05), error bars represent standard error, n=5.

**Figure 3.**
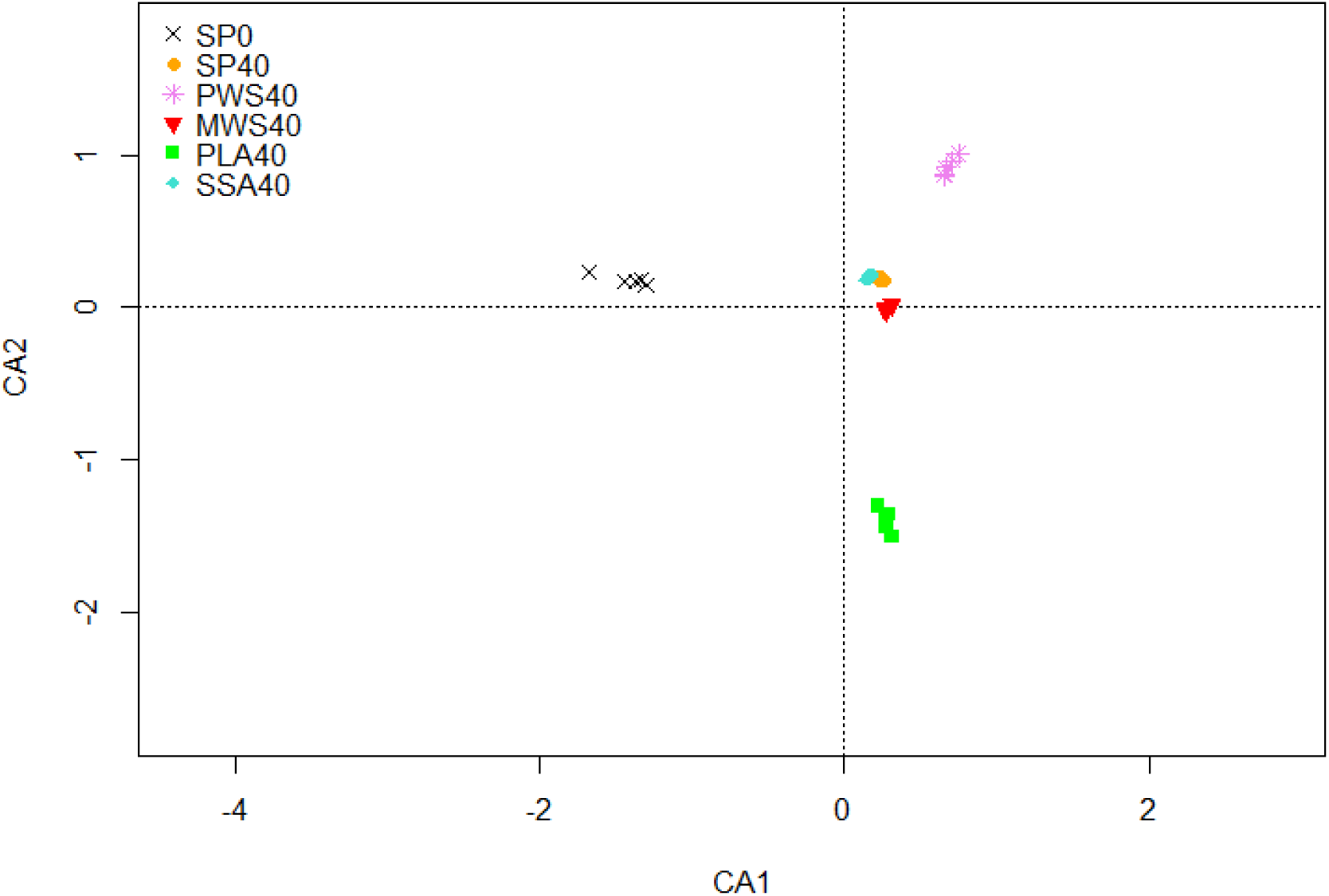
Correspondence analysis of bacterial community of the harvest after 15 months, separation between treatments was not statistically significant after pairwise comparison with Benjamini-Hochberg correction, n=5. CA1 explains 26.49% and CA2 explains 25.77% of the total variation of the data.

For the beta diversity analysis, the correspondence analysis revealed a separation of the control treatment SP0 from all other P fertilized treatments on the first axis, and a separation of PLA40 from all other treatments. (**Error! Reference source not found.**3). Overall permutational analysis was significant (P <0.001). However, after applying Benjamini-Hochberg correction of the p values, no significant differences between individual treatments were confirmed (P>0.05, Supplementary Table S2).

A CCA biplot of an ITS-DGGE abundance matrix of the harvest after 3 months showed a separation of the SP0 and SP40 treatments from all RDF treatments on the first axis, and a separation between the ash RDFs PLA40 and SSA40 from the struvite RDFs PWS40 and MWS40 on the second axis (Figure 4). Permutational analysis (999 permutations) of all treatments indicated significant differences (P<0.05), and pairwise comparison applying Benjamini-Hochberg corrections revealed significant differences between all treatments (Supplementary Table S3). The environmental factors tri-calcium phosphate solubilizing (TCP) CFUs, potential acid phosphomonoesterase (ACP) activity and plant available P measured via Morgan’s P test were significantly (P<0.05) correlated with the shift of the fungal communities.

**Figure 4.**
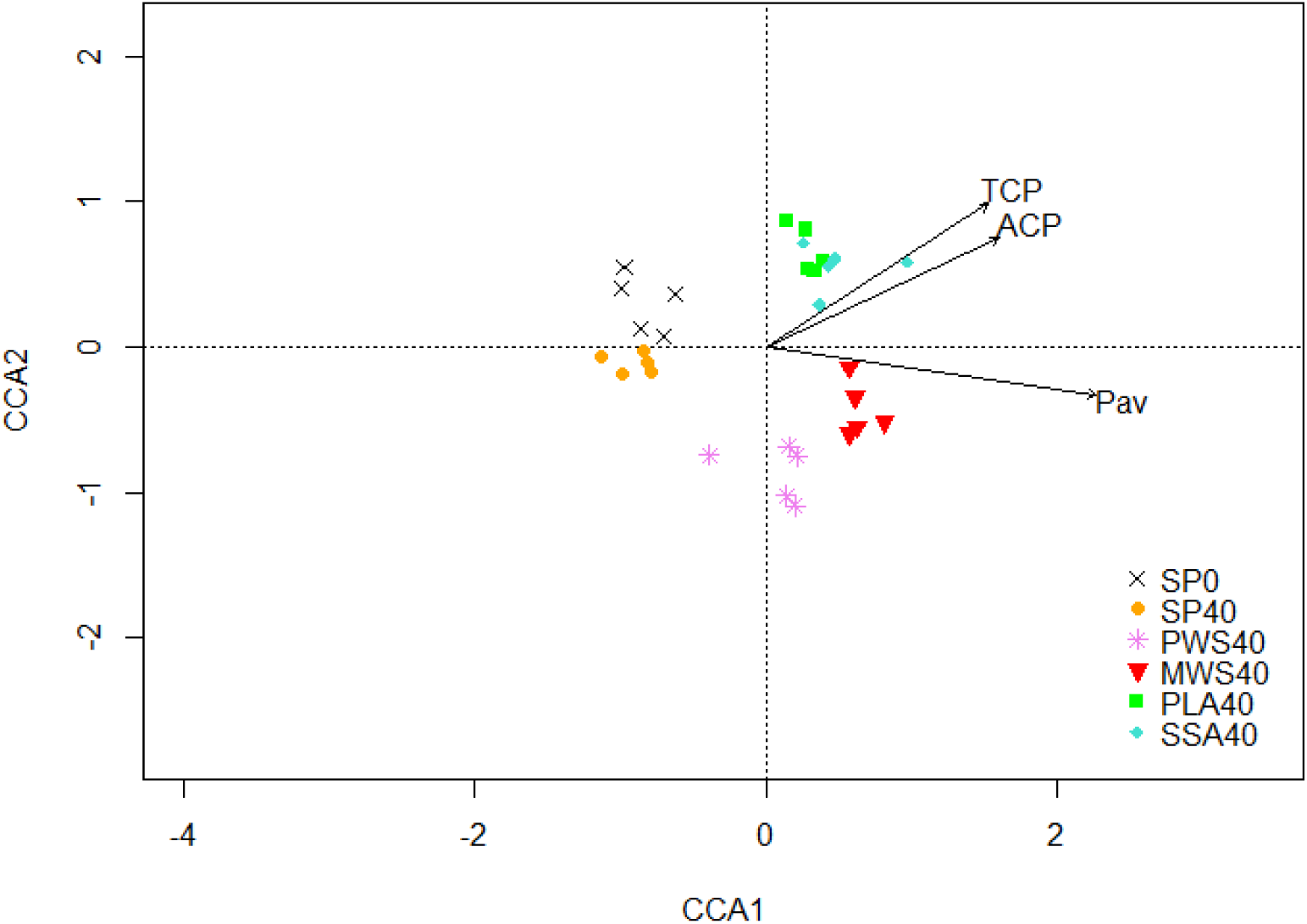
CCA biplot of ITS DGGE abundance matrix of the field trial harvest after 3 months, significant parameters involved in the shift of the fungal community are tri-calcium phosphate solubilizing bacteria (TCP), potential acid phosphomonoesterase activity (ACP) and plant available P determined via Morgan’s P test (Pav), CCA1 explains 9.94% and CCA2 7.77% of the total variation of the data, n=5.

DNA samples from the 2^nd^ year (after 15 months) were subjected to sequencing of the ITS region and 3,159,355 sequences were received. On average, 105,312 sequences per sample were applied as input into the DADA2 plug-in in QIIME2, 87.9% of those passed the filter, 82.7% were then denoised and merged, and after chimera removal, around 86,451 sequences per sample (82.1%) were obtained for further analysis and taxonomy assignment. Fisher’s alpha diversity index was significantly higher (P<0.05) for the PWS40 treatment with 60.4 compared to the treatments SP0, SP40, PLA40 and SSA40 (Supplementary Table S1). All treatments were dominated by the *Ascomycota* with a relative abundance over 80%,, followed by the phyla *Mortierellomycota*, *Zoopagomycota*, *Chytridiomycota*, *Mucoromycota*, *Basidiomycota*, *Rozellomycota*, *Glomeromycota*, *Aphelidiomycota* and *Kickxellomycota* in descending order of relative abundance (Supplementary Figure S2).

At the genus level, only two dominant genera (over 20% cumulative relative abundance across all treatments) were differentially abundant. Their relative abundance per treatment is illustrated in Figure 5. *Fusarium* was significantly higher (P<0.05) abundant in MWS40, compared to the ash treatment SSA40. The genus *Myrmecridium* was significantly higher relative abundant (P<0.05) in the SP0 and SP40 treatment, compared to both ashes (PLA40 and SSA40).

**Figure 5.**
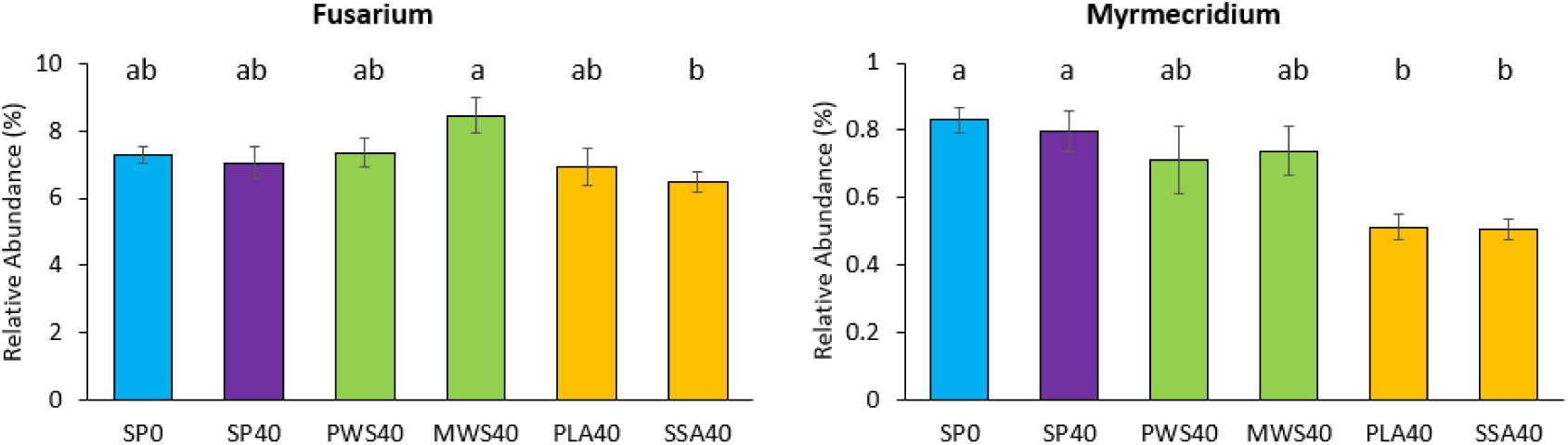
Differential abundant fungal genera (ITS sequencing) of month 15 harvest, different letters indicate significant difference (P < 0.05), error bars represent standard error, n=5.

Finally, the CA biplot of the fungal ITS community in Figure 6 shows a separation of the SP40 treatment from all others on the first axis and a separation of the SP0 control from all other treatments on the second axis. Pairwise comparison using Benjamini-Hochberg correction did not reveal any significant differences between the treatments (Supplementary Table S4). The p values without adjustment that were below 0.05 were between pairs of PWS40 and SP0, SP40 and PWS40, SP40 and PLA40, SP40 and SSA40, PWS40 and SSA40, as well as MWS40 and SSA40.

**Figure 6.**
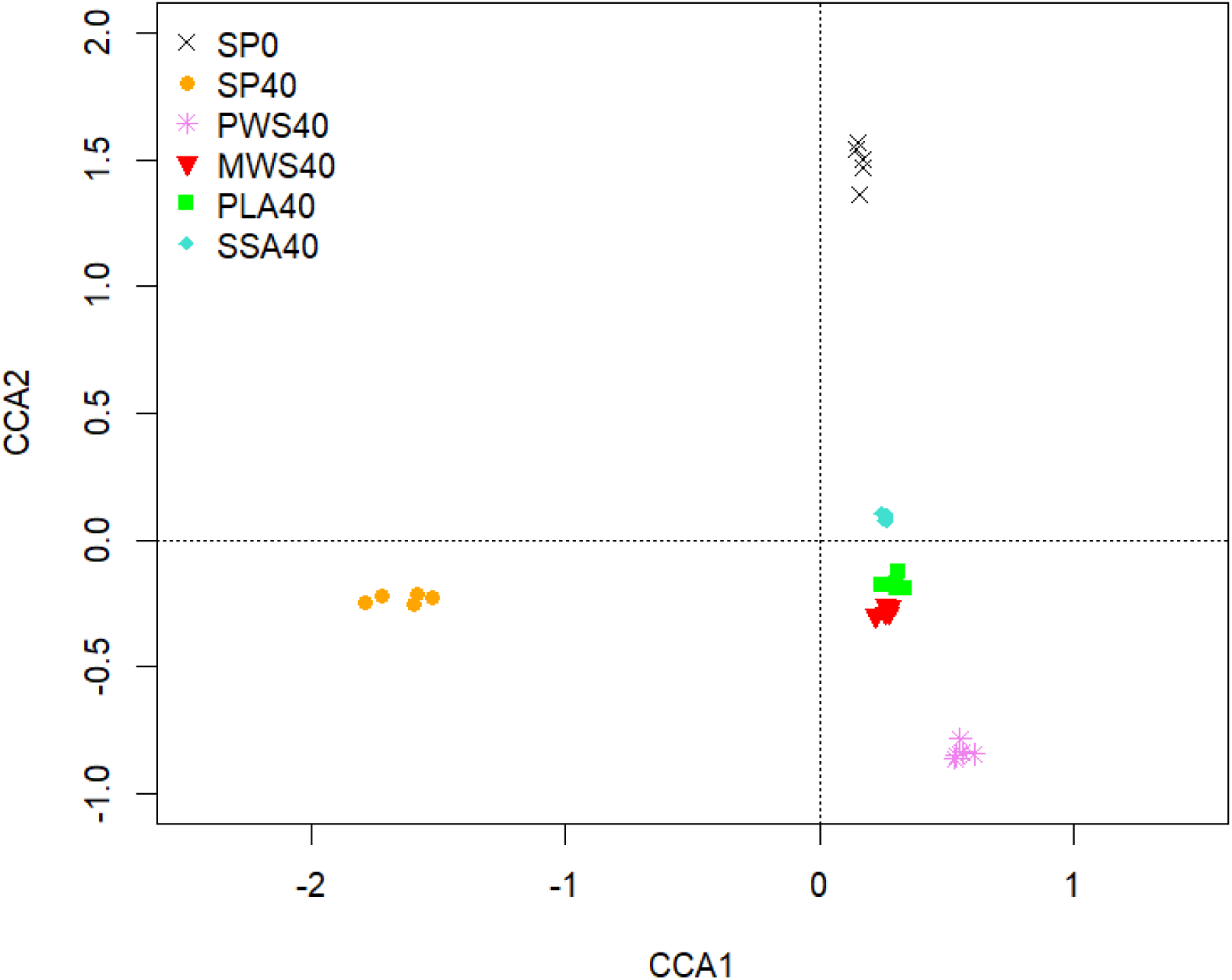
CA biplot of the ITS sequencing analysis of the harvest after 15 months, CCA1 explains 3.68% and CCA2 3.62% of the total variation of the data, n=5.

### 3.3 Analysis of Bacterial Communities Harbouring *phoD*

For the harvest after 3 months, 6,523 OTUs were picked at a similarity threshold of 75% of which 1,716,854 (80.3%) sequences were mapped to the OTUs. Alpha diversity indices ranged from 3667 to 4344 for observed features, 6.363 to 6.413 for Shannon and 0.9902 to 0.9913 for Simpson (Supplementary Table S5). Observed features were significantly different (P<0.05) between SP40 and PWS40 treatment, with PWS40 that had the higher number of observed features.

For the samples of the harvest after 15 months, 7,492 OTUs were picked at a similarity threshold of 75%, and 1,574,659 (76.3%) of the merged sequences were mapped to OTUs. Observed features ranged from 3744 to 4460, Chao1 from 4929 to 5449, ACE from 5051 to 5577, Shannon from 6.383 to 6.545 and Simpson from 0.9918 to 0.9931 (Supplementary Table S5). For the alpha diversity indices observed features, Chao1 and ACE, the SP40 treatment was in all cases significantly lower (P<0.05) than SSA40, and for observed features it was also significantly lower than PLA40. Shannon and Simpson indices were not statistically different.

Overall permutational analysis of both CAs illustrated in Figure 7 did not detect statistically significant differences in the *phoD* harbouring communities in the different P fertilized treatments. In the CA biplot for the harvest after 3 months (Figure 7A), a separation of the SSA40 treatment from all others is visible on the first axis. Pairwise comparison via PERMANOVA showed a significant difference (P<0.05) between the SP40 and PWS40 treatment, however, after applying Benjamini-Hochberg correction no significant differences were confirmed (Supplementary Table S6). In the CCA biplot of the *phoD* community of the harvest after 15 months, a similar separation of the SSA40 treatment from all other treatments is visible on the first axis, and SP0 and SP40 were separated from all other treatments on the second axis. PERMANOVA pairwise comparison of treatments showed a significant difference (P<0.05) between PWS40 and SSA40, however, after applying Benjamini-Hochberg correction, this was not confirmed.

**Figure 7.**
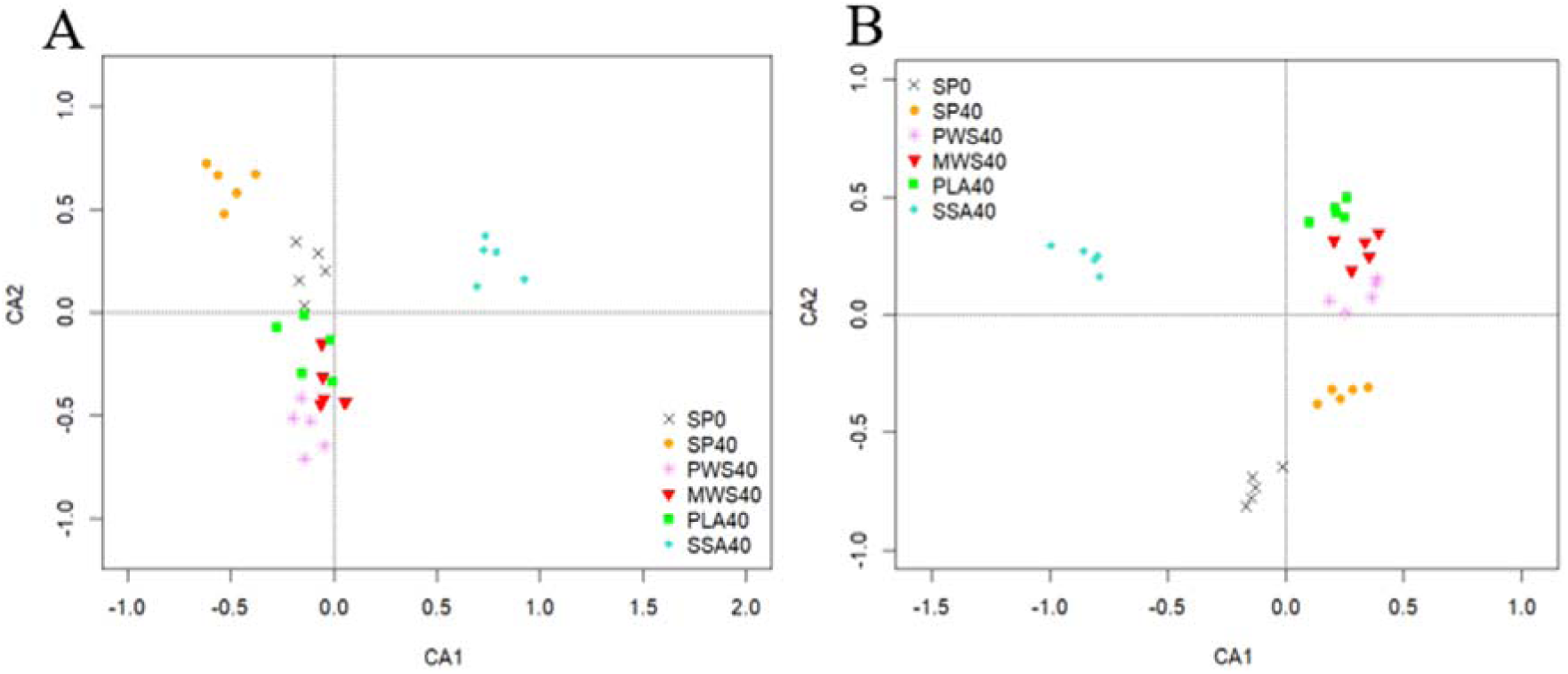
CA biplot of the phoD harbouring community A) of the harvest after 3 months, CA1 explains 4.00% and CA2 3.89% of the total variation of the data and B) of the harvest after 15 months, CA1 explains 4.14% and CA2 3.66% of the total variation of the data, n=5.

Non-parametric tests of the mean relative abundance of genera of the *phoD* sequencing analysis yielded only two genera with significantly differential relative abundance (P<0.05) for each of the two harvests (Figure 8). *Bradymonas* was significantly lower relative abundant (P<0.05) in PLA40 compared to SP0 control for the analysis of the harvest after 3 months samples, while *Actinomycetales* were significantly higher (P<0.05) relative abundant in the ash treatment SSA40 when compared to the struvite treatment PWS40 and MWS40.

**Figure 8.**
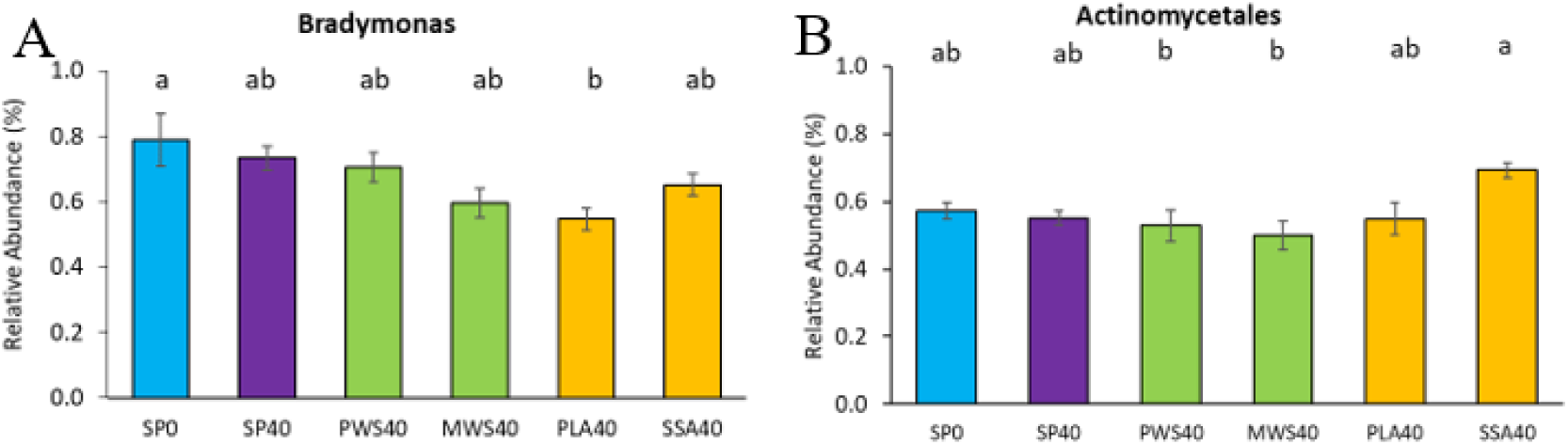
Mean relative abundance of selected genera of the phoD sequences in A) the harvest after 3 months and B) the harvest after 15 months, different letters indicate statistically significant difference (P<0.05), error bars represent standard errors, n=5.

## 4. Discussion

In this study, the impact of four RDFs on P fertilization and the soil microbial community involved in P cycling in a perennial ryegrass field trial was investigated on three time points after 3, 5 and 15 months of the start of the trial. In the beginning it was hypothesized that RDF application generates comparable plant growth promoting effects as conventional P fertilizer superphosphate in part by fostering microbial P mobilization activities.

During the first year of the field trial, TCP solubilization was enhanced for all RDF treatments, while the SP40 treatment demonstrated initially a negative impact on the abundance of mineral P solubilizing colonies. This is similar to recent findings from a short-term pot trial where three of the RDFs (one struvite, two ashes) from this study were utilized (Deinert *et al*., 2023). Many of the parameters tested in the present study (phosphonate and phytate utilization, acid and alkaline phosphomonoesterase activity and *phoC* and *phoD* gene abundance) showed strong seasonal effects that exceeded fertilizer treatment effects. While harvests 3 and 15 months after the field trial start took place in July, month 5 harvest took place in September. In September, one would expect to see the beginning of a downturn in plant production and this should be recognised. In the present study, a reduction in *phoD* copy numbers was identified in September, but the alkaline phosphatase activity remained similar to that in July. One could speculate that this drop in *phoD* copy numbers is related to an inactive sub-population of the P cycling bacteria. Further differences between months 3 and 15 were detected as the numbers of cultivable phosphonoacetic acid and phytate utilizers increased alongside the alkaline phosphatase activity, while *phoD* copy numbers were unchanged. These findings suggest that certain bacteria involved in P mobilization were more active and abundant in numbers that were not directly related to *phoD* copies.

Additionally, all cultivation dependent methods and potential ALP activity were no longer significantly different between treatments after 15 months. Seasonal effects on phosphomonoesterase activities have been reported in the past, impacting the functionality of the microbial nutrient cycle (Magid & Nielsen, 1992, Zhao *et al*., 2009). Variations in soil temperature, moisture, plant growth, root activity and organic matter accumulation have been reported before as important parameters influencing the soil P availability, mediated by microbial activity (McGrath *et al*., 2000). Only the *phoC* and *phoD* copy numbers were significantly higher in SP40 after 15 months, however this did not translate into similar findings for the potential ACP and ALP activity. Only the SSA40 treatment demonstrated significantly higher ACP activity. However, gene copy numbers are not directly correlated with enzyme activity in the soil as the sole presence of a functional gene does not indicate its translation into the relevant protein at the time of investigation. They represent the potential of alkaline and acid phosphatase activity. Furthermore, *phoD* and *phoC* are not the sole phosphatase genes that can be found among microbes nor is it likely that the chosen *phoD* and *phoC* primers will capture all variations of these marker genes (Bergkemper *et al*., 2016). Transcriptomic approaches would need to be employed to capture gene expression of the likes of *phoD* and *phoC* (Nannipieri *et al*. 2012). Furthermore, unaffected phosphomonoesterase activity does not necessarily mean that the soil microbiota remained unaffected by the treatment, the P compound hydrolysis activity could have also been compensated by other mineralization processes in soil or compensated by the presence of extracellular phosphomonoesterases only (Nannipieri *et al*. 2011).

A paucity of studies have evaluated the microbial response to novel recycling-derived fertilizer application such as struvites and ashes. Karpinska and colleagues (2021) analysed the same field trial that the present study has analysed, but their focus was exclusively on the short term-effect (three months after the trial start) on the bacterial, fungal and nematode community structure. They found that three out of the four RDFs had the same alpha diversity as the superphosphate treatment. Only SSA40 had a significantly higher Simpson’s index when compared to SP40. In this current study, the analysis of the bacterial community in the second year of the field trial (after 15 months) did not reveal any strong responses to P application. However, a significant higher abundance of *Actinobacteriota* was detected in MWS40 and SSA40 after 15 months. While trends of increased *Actinobacteriota* in P fertilized plots was also reported after 3 months, this was not statistically significant (Karpinska *et al*., 2021).

While a recent study of the field trial was not able to detected significant differences in fungal beta diversities using a NGS approach (Karpinska *et al*., 2021), analysis of the fungal community in this study based on PCR-DGGE revealed a significant separation between all treatments in the first year of the trial. Plant available P was correlated with the struvite RDFs, while tri-calcium phosphate solubilizing capabilities and acid phosphomonoesterase activity were correlated with the ash RDFs. Both ashes used in the present study were found to have low heavy metal but high metal contents. The latter may negatively influence the P availability, thus one may speculate that this reduced availability when compared to struvites may result in relatively higher microbial driven mineralization of organic P as well as immobilized inorganic P. High P availability of struvites and thus the plant growth promotion of these novel fertilizers had been reported before (Plaza *et al*., 2007, González-Ponce *et al*., 2009, Hertzberger *et al*., 2020), however little research has been conducted on the microbial response to struvite application. An increase in Actinobacterial orders was recently detected after struvite application in a *Hordeum vulgare* pot trial (Bastida *et al*., 2019). Similarly in the present study, the phylum *Actinobacteriota* was significantly higher abundant after 15 months in the struvite treatment MWS40 when compared to SP40. Observations of phosphomonoesterase activity and tri-calcium phosphate solubilizing capability in soil upon ash fertilizer application are rarely reported. Nevertheless, a study described the impact of sewage sludge ash-granule application at different rates compared to an unamended control on phosphatase activity (Możdżer, 2022). They found an increase in enzyme activity in the sludge-ash granule treatments compared to the control, however, no differentiation between acid or alkaline phosphomonoesterase activity was made and a conventional P fertilizer comparison was absent.

In the present study, the fertilizer treatment effect on the fungal communities as detected after 3 months via PCR-DGGE was much less pronounced after 15 months using fungal sequencing analysis. While it is possible that these different observations are based on fungal primer selection (which is different between DGGE and NGS sequencing), it is possible that the fertilizer event has been primarily a temporary disturbance for the fungal communities, since P fertilization took place only once (other nutrients were reapplied). It is important to note that farmers would most likely re-apply P fertilizer at least once a year if not twice and thus the more pronounced short-term effects identified in the present study could get compounded over time.

Although no significant overall effect of the fertilizer application was detected in the beta diversity of the bacterial and fungal communities after 15 months, individual phylogenetic groups were affected by the treatments. The ash treatments demonstrated some distinguishing results across the bacterial and fungal community in the present study. PLA40 had significantly higher relative abundance of *Firmicutes* and *Rokubacteria*, however the relative abundance of *Actinobacteriota* was negatively affected. SSA40 had significantly low abundances in the genera *Bacillus* and *Bradyrhizobium* (after 15 months), both well known for their involvement in P cycling activities (Halder *et al*., 1990, Khan *et al*., 2009, Zhang *et al*., 2021). In contrast, PLA40 and SSA40 both had significantly lower abundance of the cellulose-degrading fungus *Myrmecridium* that is associated with phytopathogenic activity (Marin-Felix *et al*., 2019). The genus *Fusarium* was significantly lower abundant in SSA40 as well in this study. For this genus mycotoxin production has been observed in cereals, which causes food and feed contamination worldwide, causing acute as well as chronic toxic effects in humans and animals (Ji *et al*., 2019). The struvite RDFs either displayed positive effects on the microbial P community involved in P cycling, determined via higher relative abundance of *Actinobacteriota* and *Bacillus* in MWS40, in this current study or did not show any differentiating effects at all.

## 5. Conclusion

The single application of RDFs (struvite and ash products) as a source of alternative P fertiliser in Irish grassland soil did not significantly shift the overall bacterial communities 15 months after amendments took place. Certain phyla and genera associated with P cycling activity were both positively as well as negatively affected by ash RDFs, while struvite RDFs showed positive effects on the relative abundance of P mobilizing microbes or no change at all. While the present study could not find strong positive RDF effects on the bacterial communities as initially anticipated, the lack of negative feedbacks indicates their suitability as a P fertilizer at the ecosystem level. The diversity of *phoD* was not affected by any of the fertilization treatments after 15 months, suggesting that the P mobilizing bacterial community is relatively stable in the analysed trial. However, one may speculate that repeated applications of mineral P fertilizer may have negative long-term effects and thus future long-term studies of struvites and ashes as P fertilizers with repeated applications alongside conventional superphosphate may provide clearer insights for future agriculture. RDFs showed some degree of influence on the fungal community after 3 months of application and to a lesser extend after 15 months. Likewise, SP0 and SP40 were not significantly different to the fungal community structure analysis overall. A potentially lower relative abundance of phytotoxic and mycotoxin producing fungi in the ash RDFs was detected within single application cycle. It might be worth to conduct further in-depth studies on RDF application over multiple seasons with repeated applications to investigate their effects on soil microbiota and P cycling processes. Overall, this study neither found strong positive nor negative RDF effects on the bacterial communities as initially anticipated and would recommend further studies to assess their suitability as a renewable P fertilizer at the different agronomic conditions considering other crop and soil type, application rates and frequency.

## 5. Acknowledgements

We would like to thank Interreg NWE for funding this study through the ReNu2Farm (NWE601) consortium and Cathal Redmond for his support in setting up and maintaining the field trial.

