## Supplementary for "The impact of struvite and ash recycling-derived fertilizers on microbial phosphorus mobilization capabilities and community structure in a *Lolium perenne* field trial"

**Supplementary information**

### A) Supplementary tables

*Supplementary Table S1. Alpha diversity of the 16S rRNA bacterial community and the ITS fungal community of the Teagasc field trial after 15 months, significance determined via Kruskal-Wallis test and Wilcoxon post-hoc analysis and Benjamini-Hochberg correction, different letters within a column indicate statistically significant difference, n=5.*

| *16S rRNA Sequencing* | | | | | | | | |
|---|---|---|---|---|---|---|---|---|
| **Treatment** | **Shannon** | | **Simpson** | | **InvSimpson** | | **Fisher** | |
| SP0 | 6.154[a] | ± 0.063 | 0.9914[a] | ± 0.0005 | 119.0[a] | ± 8.5 | 265.7[a] | ± 18.1 |
| SP40 | 6.054[ab] | ± 0.009 | 0.9912[a] | ± 0.0001 | 113.8[a] | ± 1.8 | 228.4[ab] | ± 4.3 |
| PWS40 | 6.037[ab] | ± 0.032 | 0.9905[a] | ± 0.0004 | 105.5[a] | ± 4.6 | 239.6[ab] | ± 6.9 |
| MWS40 | 6.056[ab] | ± 0.015 | 0.9907[a] | ± 0.0002 | 107.4[a] | ± 2.7 | 237.9[ab] | ± 3.5 |
| PLA40 | 6.051[ab] | ± 0.030 | 0.9901[a] | ± 0.0005 | 101.6[a] | ± 4.3 | 248.0[ab] | ± 6.4 |
| SSA40 | 5.911[b] | ± 0.086 | 0.9909[a] | ± 0.0003 | 110.7[a] | ± 3.9 | 196.8[b] | ± 22.5 |
| *ITS Sequencing* | | | | | | | | |
| **Treatment** | **Shannon** | | **Simpson** | | **InvSimpson** | | **Fisher** | |
| SP0 | 3.740[a] | ± 0.042 | 0.9319[a] | ± 0.0055 | 15.0[a] | ± 1.0 | 54.6[b] | ± 1.4 |
| SP40 | 3.715[a] | ± 0.058 | 0.9320[a] | ± 0.0056 | 15.0[a] | ± 1.1 | 53.6[b] | ± 1.6 |
| PWS40 | 3.760[a] | ± 0.050 | 0.9328[a] | ± 0.0042 | 15.1[a] | ± 0.8 | 60.4[a] | ± 1.2 |
| MWS40 | 3.738[a] | ± 0.057 | 0.9314[a] | ± 0.0049 | 14.9[a] | ± 1.0 | 55.1[ab] | ± 1.2 |
| PLA40 | 3.584[a] | ± 0.050 | 0.9157[a] | ± 0.0060 | 12.1[a] | ± 0.8 | 53.4[b] | ± 0.9 |
| SSA40 | 3.672[a] | ± 0.075 | 0.9239[a] | ± 0.0074 | 13.7[a] | ± 1.3 | 54.8[b] | ± 1.2 |

*Supplementary Table S2. Pairwise comparison (PERMANOVA) results of the 16S rRNA sequencing of the Teagasc field trial after 15 months, p value adjusted using the Benjamini-Hochberg procedure, statistically significant p values (P<0.05) are marked with an asterisk, n=5.*

| Treatment 1 | Treatment 2 | F | p Value | p Value adjusted |
|---|---|---|---|---|
| SP0 | SP40 | 1.227 | 0.106 | 0.179 |
| SP0 | PWS40 | 1.271 | 0.052 | 0.174 |
| SP0 | MWS40 | 1.243 | 0.141 | 0.179 |
| SP0 | PLA40 | 1.187 | 0.110 | 0.179 |
| SP0 | SSA40 | 3.712 | 0.036* | 0.174 |
| SP40 | PWS40 | 1.023 | 0.346 | 0.382 |
| SP40 | MWS40 | 1.061 | 0.247 | 0.326 |
| SP40 | PLA40 | 1.218 | 0.065 | 0.174 |
| SP40 | SSA40 | 3.196 | 0.066 | 0.174 |
| PWS40 | MWS40 | 0.962 | 0.473 | 0.535 |
| PWS40 | PLA40 | 0.898 | 0.833 | 0.841 |
| PWS40 | SSA40 | 2.993 | 0.082 | 0.174 |
| MWS40 | PLA40 | 1.009 | 0.321 | 0.382 |
| MWS40 | SSA40 | 2.981 | 0.074 | 0.174 |
| PLA40 | SSA40 | 3.280 | 0.053 | 0.174 |

*Supplementary Table S3. Pairwise comparison (PERMANOVA) results of ITS DGGE of the Teagasc field trial after 3 months, p value adjusted using the Benjamini-Hochberg procedure, statistically significant p values (P<0.05) are marked with an asterisk, n=5.*

| Treatment 1 | Treatment 2 | F | p Value | p Value adjusted |
|---|---|---|---|---|
| SP0 | SP40 | 1.821 | 0.047* | 0.046* |
| SP0 | PWS40 | 4.081 | 0.007* | 0.015* |
| SP0 | MWS40 | 5.237 | 0.007* | 0.015* |
| SP0 | PLA40 | 3.378 | 0.008* | 0.015* |
| SP0 | SSA40 | 3.673 | 0.013* | 0.015* |
| SP40 | PWS40 | 5.735 | 0.006* | 0.015* |
| SP40 | MWS40 | 7.538 | 0.012* | 0.015* |
| SP40 | PLA40 | 6.004 | 0.013* | 0.015* |
| SP40 | SSA40 | 6.140 | 0.007* | 0.015* |
| PWS40 | MWS40 | 3.770 | 0.009* | 0.015* |
| PWS40 | PLA40 | 5.348 | 0.006* | 0.015* |
| PWS40 | SSA40 | 4.374 | 0.006* | 0.015* |
| MWS40 | PLA40 | 3.859 | 0.007* | 0.015* |
| MWS40 | SSA40 | 3.264 | 0.011* | 0.015* |
| PLA40 | SSA40 | 2.770 | 0.029* | 0.031* |

*Supplementary Table S4. Pairwise comparison using PERMANOVA of the fungal ITS sequencing of the Teagasc field trial after 15 months, p value adjusted using the Benjamini-Hochberg procedure, statistically significant p values (P<0.05) are marked with an asterisk, n=5.*

| Treatment 1 | Treatment 2 | F | p Value | p Value adjusted |
|---|---|---|---|---|
| SP0 | SP40 | 0.980 | 0.604 | 0.626 |
| SP0 | PWS40 | 1.147 | 0.023* | 0.087 |
| SP0 | MWS40 | 1.074 | 0.118 | 0.153 |
| SP0 | PLA40 | 1.098 | 0.102 | 0.145 |
| SP0 | SSA40 | 1.136 | 0.074 | 0.145 |
| SP40 | PWS40 | 1.178 | 0.009* | 0.060 |
| SP40 | MWS40 | 1.083 | 0.107 | 0.153 |
| SP40 | PLA40 | 1.190 | 0.008* | 0.060 |
| SP40 | SSA40 | 1.201 | 0.047* | 0.138 |
| PWS40 | MWS40 | 0.990 | 0.524 | 0.549 |
| PWS40 | PLA40 | 1.045 | 0.231 | 0.310 |
| PWS40 | SSA40 | 1.253 | 0.006* | 0.060 |
| MWS40 | PLA40 | 1.107 | 0.078 | 0.145 |
| MWS40 | SSA40 | 1.298 | 0.017* | 0.060 |
| PLA40 | SSA40 | 1.025 | 0.414 | 0.410 |

*Supplementary Table S5. Alpha diversity of phoD functional gene sequencing results of the Teagasc field trial after 3 and 15 months, significance determined via Kruskal-Wallis test and Wilcoxon post-hoc analysis and Benjamini-Hochberg correction, different letters within a column indicate statistically significant difference, n=5.*

| **Treatment** | **Observed Features** | | **Chao1** | | **ACE** | | **Shannon** | | **Simpson** | |
|---|---|---|---|---|---|---|---|---|---|---|
| | | | | | *Harvest after 3 Months* | | | | | |
| SP0 | 3910.0[ab] | ± 98.18 | 4810.1[a] | ± 54.06 | 4901.7[a] | ± 44.38 | 6.370[a] | ± 0.0458 | 0.9907[a] | ± 0.0008 |
| SP40 | 3667.4[b] | ± 138.42 | 4726.5[a] | ± 124.76 | 4803.3[a] | ± 114.92 | 6.413[a] | ± 0.0443 | 0.9913[a] | ± 0.0004 |
| PWS40 | 4344.0[a] | ± 93.40 | 5008.1[a] | ± 58.75 | 5038.9[a] | ± 51.71 | 6.363[a] | ± 0.0496 | 0.9902[a] | ± 0.0007 |
| MWS40 | 3843.8[ab] | ± 206.46 | 4750.9[a] | ± 141.47 | 4812.1[a] | ± 124.97 | 6.366[a] | ± 0.0519 | 0.9908[a] | ± 0.0006 |
| PLA40 | 3887.6[ab] | ± 132.67 | 4844.5[a] | ± 101.52 | 4899.7[a] | ± 97.76 | 6.413[a] | ± 0.0714 | 0.9911[a] | ± 0.0010 |
| SSA40 | 3916.4[ab] | ± 94.43 | 4810.3[a] | ± 73.45 | 4874.9[a] | ± 64.70 | 6.406[a] | ± 0.0412 | 0.9910[a] | ± 0.0005 |
| | | | | | *Harvest after 15 Months* | | | | | |
| SP0 | 4248.8[ab] | ± 138.05 | 5260.7[ab] | ± 100.65 | 5357.5[ab] | ± 96.04 | 6.543[a] | ± 0.0326 | 0.9931[a] | ± 0.0002 |
| SP40 | 4109.0[ab] | ± 195.94 | 5207.6[ab] | ± 117.26 | 5344.2[ab] | ± 102.67 | 6.500[a] | ± 0.0520 | 0.9926[a] | ± 0.0005 |
| PWS40 | 3744.6[b] | ± 94.81 | 4929.3[b] | ± 78.62 | 5051.3[b] | ± 73.57 | 6.383[a] | ± 0.0501 | 0.9918[a] | ± 0.0006 |
| MWS40 | 4100.4[ab] | ± 142.13 | 5157.9[ab] | ± 130.69 | 5217.5[ab] | ± 129.73 | 6.477[a] | ± 0.0590 | 0.9928[a] | ± 0.0006 |
| PLA40 | 4448.4[a] | ± 74.35 | 5352.7[ab] | ± 67.09 | 5434.4[ab] | ± 56.60 | 6.482[a] | ± 0.0531 | 0.9925[a] | ± 0.0006 |
| SSA40 | 4460.8[a] | ± 153.76 | 5449.0[a] | ± 93.05 | 5576.6[a] | ± 77.14 | 6.545[a] | ± 0.0351 | 0.9931[a] | ± 0.0002 |

*Supplementary Table S6. Pairwise comparison using PERMANOVA of the phoD sequencing of the harvest after 3 and 15 months of the Teagasc field trial, p value adjusted using the Benjamini-Hochberg procedure, statistically significant p values (P<0.05) are marked with an asterisk, n=5.*

| *Harvest after 3 Months* | | | | |
|---|---|---|---|---|
| **Treatment 1** | **Treatment 2** | **F** | **p Value** | **p Value adjusted** |
| SP0 | SP40 | 1.021 | 0.394 | 0.681 |
| SP0 | PWS40 | 0.885 | 0.866 | 0.936 |
| SP0 | MWS40 | 0.947 | 0.560 | 0.805 |
| SP0 | PLA40 | 0.971 | 0.449 | 0.713 |
| SP0 | SSA40 | 0.992 | 0.435 | 0.713 |
| SP40 | PWS40 | 1.167 | 0.150 | 0.681 |
| SP40 | MWS40 | 1.143 | 0.210 | 0.681 |
| SP40 | PLA40 | 1.039 | 0.305 | 0.681 |
| SP40 | SSA40 | 1.316 | 0.015* | 0.270 |
| PWS40 | MWS40 | 0.851 | 0.876 | 0.936 |
| PWS40 | PLA40 | 0.850 | 0.894 | 0.936 |
| PWS40 | SSA40 | 1.094 | 0.212 | 0.681 |
| MWS40 | PLA40 | 0.811 | 0.942 | 0.936 |
| MWS40 | SSA40 | 1.021 | 0.340 | 0.681 |
| PLA40 | SSA40 | 1.040 | 0.353 | 0.681 |
| *Harvest after 15 Months* | | | | |
| **Treatment 1** | **Treatment 2** | **F** | **p Value** | **p Value adjusted** |
| SP0 | SP40 | 0.996 | 0.444 | 0.494 |
| SP0 | PWS40 | 1.165 | 0.063 | 0.405 |
| SP0 | MWS40 | 1.029 | 0.412 | 0.494 |
| SP0 | PLA40 | 1.052 | 0.271 | 0.494 |
| SP0 | SSA40 | 1.005 | 0.417 | 0.494 |
| SP40 | PWS40 | 1.011 | 0.408 | 0.494 |
| SP40 | MWS40 | 1.054 | 0.287 | 0.494 |
| SP40 | PLA40 | 1.051 | 0.314 | 0.494 |
| SP40 | SSA40 | 1.197 | 0.081 | 0.405 |
| PWS40 | MWS40 | 0.980 | 0.534 | 0.574 |
| PWS40 | PLA40 | 1.058 | 0.238 | 0.494 |
| PWS40 | SSA40 | 1.321 | 0.038* | 0.405 |
| MWS40 | PLA40 | 0.840 | 0.891 | 0.906 |
| MWS40 | SSA40 | 1.201 | 0.089 | 0.405 |
| PLA40 | SSA40 | 1.073 | 0.257 | 0.494 |

**B) Supplementary figures**

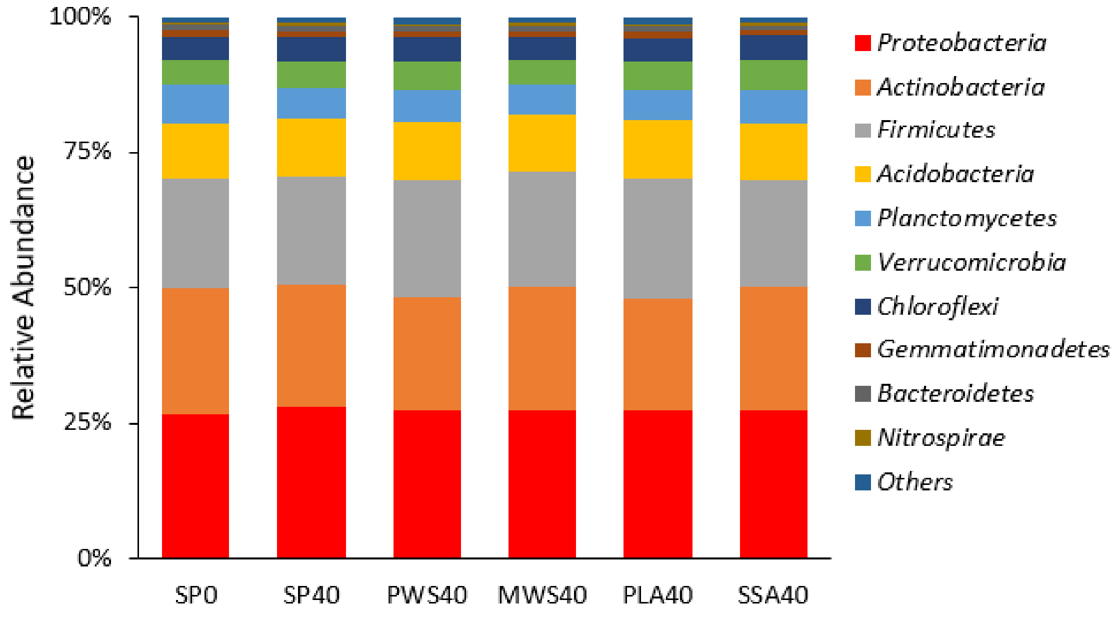


*Supplementary Figure S1. Mean relative abundance (%) depicted as stacked bar plot of the top ten relative abundant phyla identified in the 16S rRNA sequencing analysis of the Teagasc field trial after 15 months, all other, lower abundant phyla are combined in "Others".*

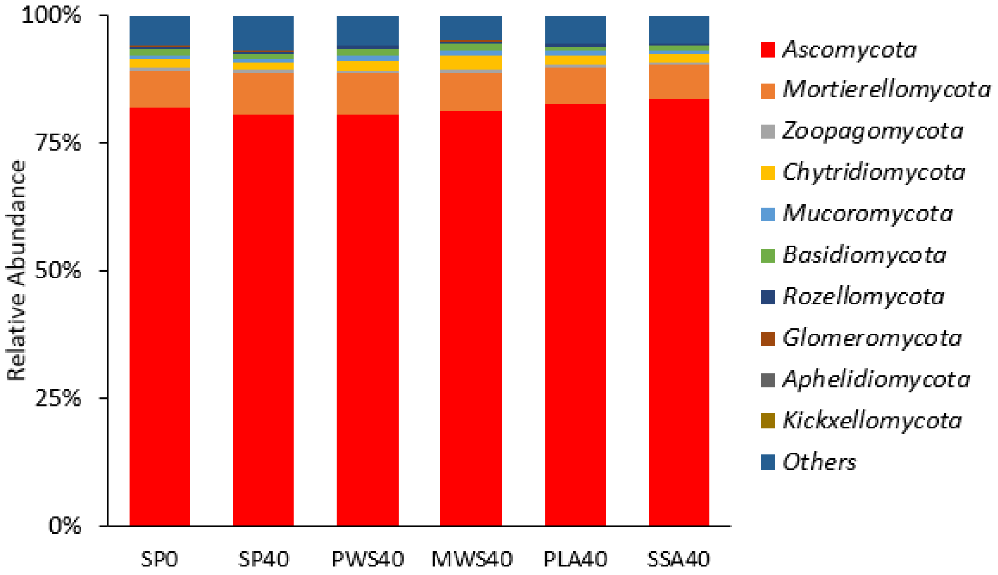


*Supplementary Figure S2. Mean relative abundance (%) of the top ten abundant phyla of the ITS sequencing analysis of the harvest after 15 months of the Teagasc field trial, n=5.*
